# The wisdom of stalemates: consensus and clustering as filtering mechanisms for improving collective accuracy

**DOI:** 10.1101/2020.01.09.899054

**Authors:** Claudia Winklmayr, Albert B. Kao, Joseph B. Bak-Coleman, Pawel Romanczuk

**Affiliations:** Bernstein Center for Computational Neuroscience, Berlin, Germany; Max-Planck-Institut für Mathematik in den Naturwissenschaften, Leipzig, Germany; Santa Fe Institute, Santa Fe, NM, USA; Department of Ecology and Evolutionary Biology, Princeton University, Princeton, NJ, USA; Institute for Theoretical Biology, Department of Biology, Humboldt Universität zu Berlin, Germany

**Author notes:** These authors contributed equally to this work (Claudia Winklmayr, Albert B. Kao).

**Keywords:** collective decision making, collective intelligence, consensus decision, voter model, wisdom of crowds

## Abstract

Groups of organisms, from bacteria to fish schools to human societies, depend on their ability to make accurate decisions in an uncertain world. Most models of collective decision-making assume that groups reach a consensus during a decision-making bout, often through simple majority rule. In many natural and sociological systems, however, groups may fail to reach consensus, resulting in stalemates. Here, we build on opinion dynamics and collective wisdom models to examine how stalemates may affect the wisdom of crowds. For simple environments, where individuals have access to independent sources of information, we find that stalemates improve collective accuracy by selectively filtering out incorrect decisions. In complex environments, where individuals have access to both shared and independent information, this effect is even more pronounced, restoring the wisdom of crowds in regions of parameter space where large groups perform poorly when making decisions using majority rule. We identify network properties that tune the system between consensus and accuracy, providing mechanisms by which animals, or evolution, could dynamically adjust the collective decision-making process in response to the reward structure of the possible outcomes. Overall, these results highlight the adaptive potential of *stalemale filtering* for improving the decision-making abilities of group-living animals.

## Introduction

Collective decision-making is an essential feature for organisms across a wide range of taxa, from bacteria to fish to humans [1]. For some species, individuals accrue benefits from group-living for reasons unrelated to the decision-making process and make consensus decisions simply to maintain cohesiveness [2, 3]. Beyond cohesion, many other species make decisions collectively in ways that improve accuracy and the fitness of the animals within the group [4, 5].

The potential for decision accuracy to increase with group size is often referred to as the ‘wisdom of crowds,’‘collective wisdom,’ or ‘collective intelligence.’ Traditional models assume that the increase in accuracy occurs because individuals contribute different, and somewhat uncorrelated, pieces of information relevant to the decision. Because the group as a whole has access to a greater amount of information than any single individual, the resulting collective decision has the potential to be more accurate than is possible for an individual. A wide variety of theoretical models (*e.g*., [6, 7, 8]), including the well-known Condorcet jury theorem [9], and an increasing number of empirical studies (*e.g*., [10, 11]), have indeed demonstrated this effect for different contexts and species.

Models of collective wisdom often explicitly assume decision-making processes inherently lead to consensus, typically as a result of a voting process such as majority rule, quorums [12, 13, 14], or an averaging of individual opinions [6]. Realistic decision-making in animal groups, however, is not exogenously aggregated and instead relies on endogenous processes of typically local social interactions. For example, in fish schools and bird flocks, the trajectories of individual animals are influenced by both the movements of their near neighbors, as well as their own preferences, and these myriad momentary interactions may result in coherent collective movement towards a single direction of motion [15, 16, 17, 18]. While these dynamics may at times approximate majority rule [12, 13, 19], the mapping between the microscopic social interactions and macroscopic collective decisions remains an active area of research.

Crucially, emergent decision-making processes through social interactions may not guarantee that a group reaches a consensus within a reasonable time period, or at all. For example, many small schooling fish are prey to larger predators and prefer to hide in grasses or shelters for safety [20]. As a result, they often must make decisions about when to leave their shelter and where to travel (*e.g*., to acquire food). These decisions may depend on individually-sensed information about foraging opportunities and the likelihood of encountering a predator. Failure to reach a consensus, for example for one of the available food patches, and consequently staying in the shelter, may be a relatively low-cost option (compared to being eaten). The decision-making process for these species, then, may be more accurately described as a decision whether to leave the shelter or not, as well as a decision of which option (*e.g*., food patch) to visit if leaving the shelter.

Furthermore, recent work has suggested that the specific informational environment in which collective decisions are made can have major effects on the resulting decision accuracy. In particular, a combination of both uncorrelated and correlated information can interact, resulting in low decision accuracy for large group sizes, with decision accuracy instead being maximized by intermediate-sized groups [19, 21, 22]. This stands in contrast to the predictions of many wisdom of crowds models, including the Condorcet jury theorem, which predict a monotonic increase in collective accuracy as group size increases.

The presence of both uncorrelated and correlated information may frequently occur in nature due to differences in the degree of spatial and temporal correlation of different environmental cues. For example, loud auditory cues may be highly correlated across individuals in a group, while visual cues may be more localized and therefore less correlated across individuals due to the limited visual field of each animal, and occlusion of one’s field of view by the bodies of neighbors. Another way in which uncorrelated and correlated information can be present simultaneously is from social copying within a group. A theoretical model has demonstrated that if individuals begin with independent (uncorrelated) information, but make individual decisions by incorporating the previous decisions of other group mates, then a similar phenomenon of an optimal intermediate group size, and poor decision accuracy for large groups, can also emerge [23].

Opinion dynamics models provide a flexible method to examine the mapping between social interactions, environmental cues, collective decisions, and decision accuracy [24]. In such models, each individual begins with some initial opinion (e.g., determined by the environmental cues that the individual observes), and then the opinions may change in time depending on the social network structure and the social interaction rules that individuals follow. These dynamics may lead to consensus, whereby all individuals have the same opinion, or may fail to reach consensus, by arriving at some other state where the system indefinitely retains a mixture of opinions.

In order to better understand how animals may make collective decisions in naturalistic conditions, here we examine the effect of opinion dynamics on collective decisions in both simple environments (*i.e*., individuals independently sample a single environmental cue, identical to the context described by the Condorcet jury theorem) and complex environments (i.e, individuals can sample from both a correlated and an uncorrelated environmental cue). We show that opinion dynamics can substantially alter the outcome of collective decisions, and in most cases, improve collective accuracy compared to simple majority rule.

## Results

### Stalemates can improve the Condorcet jury theorem

Individuals in a group are faced with two options (e.g., two potential food patches, fleeing directions, locations to rest, etc.). If the group fails to come to a consensus on which option to choose, it effectively chooses a third option: to do nothing. Throughout this work we will assume that groups are faced with binary decisions between a ‘correct’ and an ‘incorrect’ option, and we calculate collective accuracy by taking the trial-average across those trials where consensus was reached.

To set the initial opinions of each individual, we assume, as in the Condorcet jury theorem, that each individual has access to an independently sampled environmental cue. As a result, the initial opinions are independent of each other, and each has a probability *r* of being correct (and a probability 1 − *r* of being incorrect). We assume that *r* > 1/2 (*i.e*., that the cue is positively informative, otherwise individuals could simply reverse their interpretation of the cue to generate an informative cue). As is well-known from the Condorcet jury theorem and related work, if *r* > 1/2, and a group makes decisions using simple majority rule, then the probability that the group makes a correct decision increases monotonically with group size and asymptotes at perfect accuracy (the so-called ‘wisdom of crowds’) [6].

In our model, individuals in a group are connected to one another into a social network, *i.e*., a graph where the nodes represent the individuals and connections between individuals are represented as edges. Any social network structure in principle may be used, but here we use the Watts-Strogatz network, a well-studied family of networks that spans the range from highly-clustered networks to random networks [25, 26]. Watts-Strogatz networks are particularly convenient because they are completely determined by only three parameters: The group size (*N*), the node degree (i.e., the number of neighbors that an individual is connected to, *k*) and the rewiring probability (*β*). The rewiring probability specifies the probability, when constructing the network, that a node is connected to a randomly selected node instead of a spatially-close node, thus controlling the extent to which the network is highly clustered (*β* = 0) or random (*β* = 1). Once a network has been generated, it remains static for the duration of the decision-making process. We examine groups ranging in size from *N* = 3 to 200, and, for simplicity, initially fix the number of neighbors to *k* = 5 (and a fully connected network if *N* < *k* + 1) for each individual (which approximates the number of neighbors that animals tend to pay attention to when making collective decisions [27]) and the rewiring probability to *β* = 0.2. However, we later explicitly examine the role that *k* and *β* play in affecting collective accuracy.

Next, the social interaction rules must be specified, which govern how individuals change their opinion due to the opinions of others. To approximate the social interactions among real animals, we assumed an asynchronous updating policy where at each time step, a random focal individual is selected, and that individual changes its opinion to the majority opinion of its neighbors (Figure 1A) (in the case of a tie, the individual’s opinion is left unchanged, Figure 1B) [28]. Such ‘threshold voter model’ dynamics are not guaranteed to reach consensus; instead, the network may become ‘stuck’ in an intermediate state or oscillate between intermediate states (a detailed discussion of the convergence behavior of the threshold model can be found in [29], and a minimal example of a stuck network is shown in Figure 1C). Because of this possibility, we add a further parameter, *s_max_*, which specifies the maximum number of opinion updates before ending the simulation if consensus has not been reached. We set *s_max_* to be at least twice as large as the average number of steps needed for consensus so that simulations are typically not incorrectly classified as having reached a stalemate.

**Figure 1.**
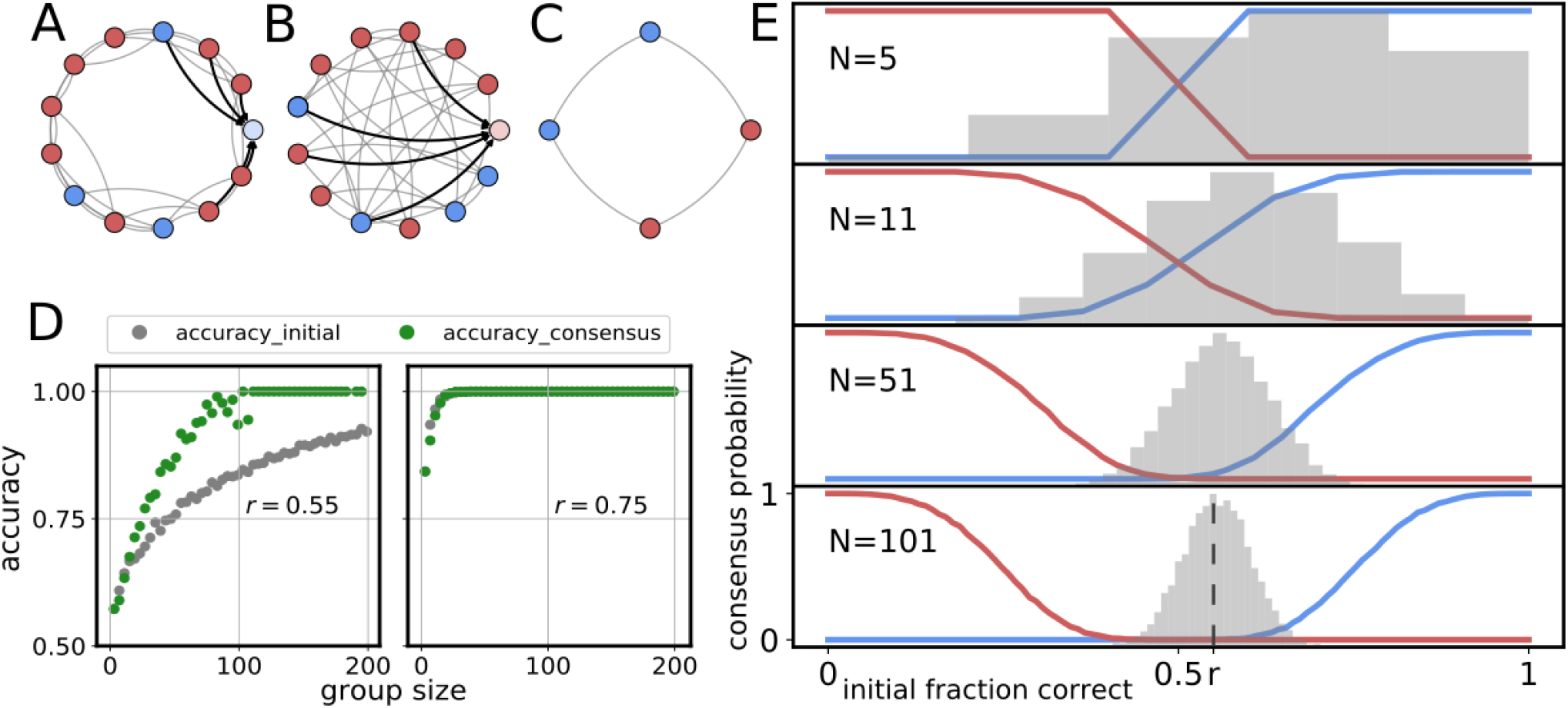
Opinion dynamics improve collective decision accuracy in simple environments. **A.** Example of a single updating step in a highly clustered Watts-Strogatz network (*β* = 0.1). The focal node (light blue) observes the opinions of *k* = 5 immediate neighbors and will change its opinion from blue to red in order to match the majority of its neighbors. **B.** Same as A but with a highly random network (*β* ≈ 1). Here the focal node (light red) observes the opinions of *k* = 4 neighbors but will maintain its opinion because its neighbors have equal numbers of red and blue opinions. **C.** A minimal example of a network that has reached a stalemate. Because each node has one blue and one red neighbor, none of the nodes will change their opinion. **D.** Examples of the adaptiveness of opinion dynamics for two different values of the cue reliability. When the cue reliability is low (*r* =0.55, left), we find that groups that reach consensus through opinion dynamics (green) show higher collective accuracy than groups which use simple majority vote (grey). When the cue reliability is high (*r* =0.75, right) collective accuracy is similar, whether a consensus is reached through opinion dynamics or simple majority rule. **E.** The probability that a group reaches a consensus for the correct (blue line) or incorrect (red line) option as a function of the initial fraction of individuals voting for the correct option. The four panels show the results for different group sizes (*N* = 5 through 101 from top to bottom). The grey histograms illustrate the distribution of initial votes for a cue with reliability *r* = 0.55. As group size increases, the initial vote distribution needs to be increasingly biased in order for a consensus to be reached (*i.e*., the inflection point of the red and blue curves shift to more extreme values). Assuming that the cue is informative (i.e., *r* > 0.5), the set of initial opinions will tend to have a positive bias, and the opinion dynamics will tend to reach consensus towards the correct option.

When networks do reach a consensus, we observe that the opinion dynamics result in an average collective accuracy that exceeds that predicted by the Condorcet jury theorem (where groups make collective decisions through simple majority rule). This is true for all values of the environmental cue reliability *r*, but particularly when the cue is unreliable (*r* ≈ 1/2) (Figure 1D). Therefore, the process of consensus formation, or, conversely, the presence of stalemates, appears to preferentially filter out inaccurate decisions and allow consensus only when the decision tends to be correct.

To illustrate the mechanism underlying this phenomenon, we plot the probability that a group makes a correct consensus decision, and the probability that group makes an incorrect consensus decision, as functions of the fraction of individuals with a correct initial opinion, for different group sizes (Figure 1E, red and blue lines). The probability of a correct (incorrect) consensus opinion increases nonlinearly with the proportion of correct (incorrect) initial opinions. We note that the probabilities of correct consensus and incorrect consensus do not necessarily sum to one because of the nonzero probability of stalemates. In particular, when there are similar numbers of correct and incorrect initial opinions (*i.e*., when the proportion of initial correct opinions is ≈ 0.5), the group is highly unlikely to reach consensus, especially at larger group sizes. Consensus reliably occurs only when the initial opinions are highly biased (*i.e*., the proportion of initial correct opinions is close to 0 or 1).

This explains the ability of stalemates to act as a filter to improve collective decisions. The probability that a certain proportion of initial opinions is correct follows a binomial distribution with mean *r* (Figure 1E, grey histograms). Since we assume that *r* > 1/2, if a particular group exhibits a highly biased set of initial opinions, it is much more likely that the initial opinions are biased in favor of the correct option than the incorrect option. Because consensus tends to occur only when the initial opinions are sufficiently biased, the function of stalemates is to reject scenarios where the majority opinion is wrong, and boost the probability of correct decisions.

### Stalemates can restore the wisdom of crowds in complex informational environments

We next investigate the effect of opinion dynamics in more informationally complex (and naturalistic) environments (Figure 2A). Here, following previous studies [19, 21, 22], we assume that there are two cues in the environment. One cue (the uncorrelated cue) is independently sampled by each individual in the group, and correctly indicates to an individual which of the two options is better with a probability *r*_I_ > 1/2. The other cue, however, is correlated across all of the individuals in the group, such that all of the individuals observe the same information from that cue in any given trial. The probability that the correlated cue correctly indicates to all individuals which of the two options is better is given by *r_C_* > 1/2. An individual in the group then sets its initial opinion to the information given by the uncorrelated cue with probability *p*, and otherwise sets its initial opinion to that given by correlated cue with probability 1 − *p*. All other aspects of the model are the same as before (*e.g*., individuals are connected to each other via a Watts-Strogatz network, and randomly selected individuals change their opinion to that of the majority of their neighbors).

**Figure 2.**
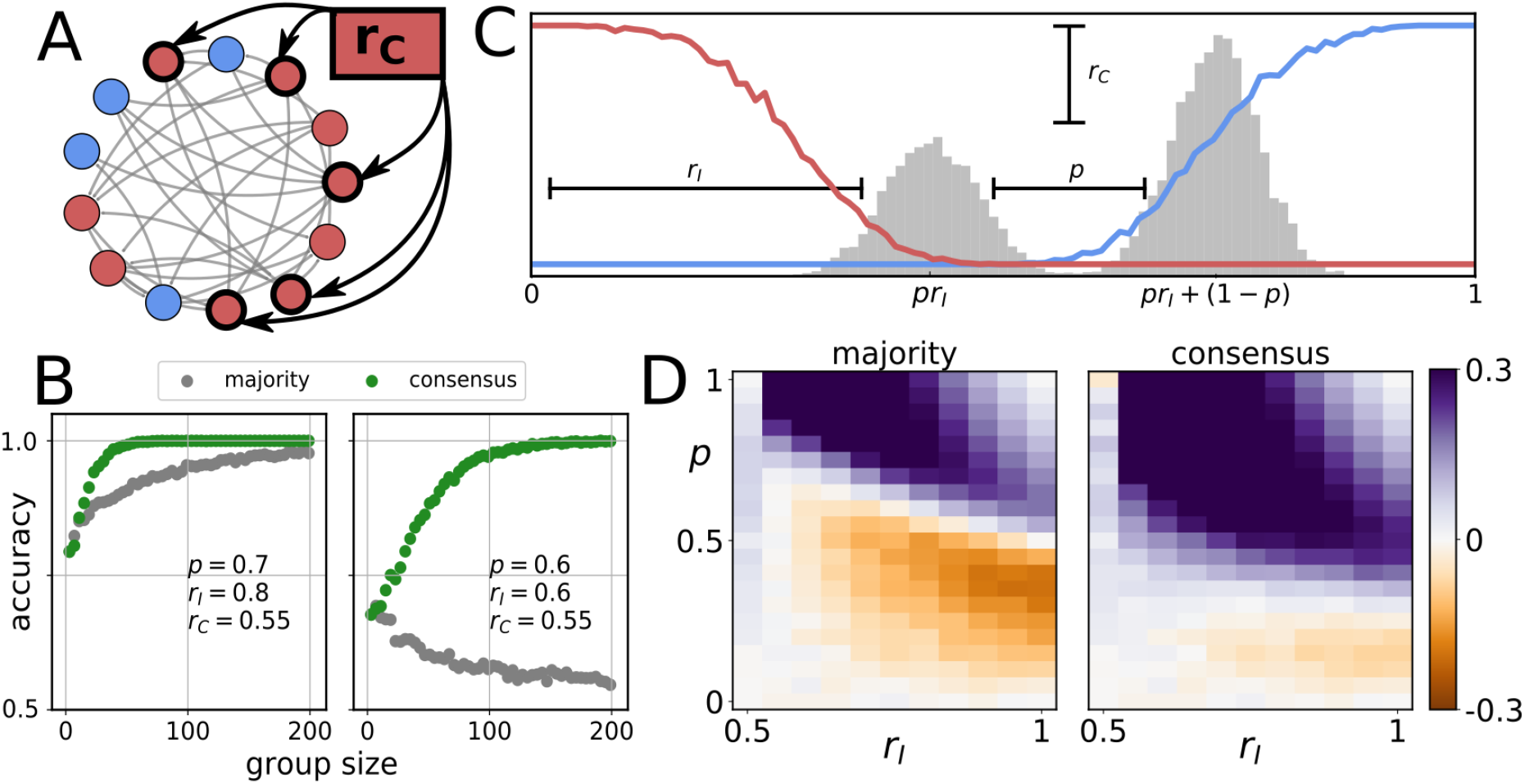
Opinion dynamics improve collective decision accuracy in complex environments. **A.** Example of the formation of initial opinions in a complex environment. With probability 1 − *p*, an individual attends to the correlated source (red box), and all individuals that follow this source will receive the same information, which is correct with probability *r_C_*. All other agents sample independently-observed information which is correct with probability *r_I_*. **B.** Opinion dynamics can improve collective accuracy. In the region of parameter space where, using simple majority rule, we observe the wisdom of crowds (*i.e*., *p* > 1/(2*r_I_*)), we observe a quantitative improvement in collective accuracy (left; *p* =0.7, *r_I_* = 0.8, *r_C_* = 0.55). However, in the region of parameter space where large groups perform poorly (*i.e*., *p* < 1/(2*r_I_*)), opinion dynamics restores the wisdom of crowds, resulting in a qualitative improvement in collective accuracy (right; *p* = 0.6, *r_I_* = 0.6, *r_C_* = 0.55). **C.** When individuals sample from one of two available cues, the distribution of initial votes is bimodal. The centers of the modes correspond to the conditional probability of an individual being correct, given that the correlated cue is correct (right mode) or incorrect (left mode). Black lines illustrate the effect of the three model parameters on the shape of the distribution: the reliability of the independent cue (*r_I_*) determines the distance from 0, the reliability of the correlated cue (*r_C_*) determines the relative heights of the two modes, and the strategy parameter (*p*) governs the distance between the modes. Red and blue lines depict the probability of a correct or incorrect consensus as in Figure 1E. Consensus is relatively unlikely when the correlated cue is incorrect (left mode). **D.** The effect of opinion dynamics in complex environments for *r_C_* = 0.55 and *N* = 51 across the entire *r_I_ × p* parameter space. Relative collective accuracy when making decisions using simple majority rule (left), or using opinion dynamics (right), compared to a solitary individual. Here we see that the region in parameter space where groups perform better than solitary decision makers (purple regions) is larger when opinion dynamics are used.

As in the simple environments, in complex environments we observe that stalemates improve collective accuracy, when the group reaches consensus. However, in contrast with the simple environments, in complex environments we often observe substantially larger increases in collective accuracy.

Previous work demonstrated that when groups use simple majority rule to make collective decisions in such complex environments, the ‘wisdom of crowds’ (*i.e*., collective accuracy increasing monotonically with group size and asymptoting at perfect accuracy) is observed only if *p* > 1/(2*r_I_*). In this region of parameter space, similar to our observations in simple environments, when a group uses opinion dynamics to arrive at a consensus collective decision, we observe a quantitative improvement over simple majority rule (Figure 2B, left panel).

By contrast, if *p* < 1/(2*r_I_*)), using simple majority rule results in a finite optimal group size, and very large groups asymptote at an accuracy of just *r_C_*, since in this region of parameter space the correlated cue tends to dominate the collective decision [21]. Here, we observe a qualitatively different effect of using opinion dynamics to make collective decisions, with social interactions causing a much larger increase in collective accuracy (Figure 2B). Indeed, when groups use opinion dynamics to reach a consensus, we find that the wisdom of crowds can be restored, and the dominance of the correlated cue negated, with collective accuracy again increasing monotonically with group size and asymptoting at perfect accuracy (Figure 2B, right panel).

To illustrate the effect of consensus decisions more broadly, we performed a parameter scan across the entire *r_I_ × p* plane, while keeping the reliablity of the correlated cue fixed at *r_C_* = 0.55 (Figure 2D). The two panels of Figure 2D show the relative collective accuracy of a group of size *N* = 51 compared to the accuracy of a solitary individual. We find that the region in parameter space where groups perform better than solitary decision makers is larger when opinion dynamics (right panel) are used than when majority rule is employed (left panel).

The filtering mechanism responsible for this qualitative improvement in collective accuracy is similar as for simple environments: the network tends to reach a stalemate if the set of initial opinions is balanced and is increasingly likely to reach consensus if the initial opinions are more biased towards one of the two options. However, the probability distribution of initial opinions is substantially different in complex informational environments compared to simple environments (Figure 2C). In complex environments, we observe a bimodal distribution of initial opinions, one mode resulting from cases where the correlated cue is correct, and the other mode resulting from cases where the correlated cue is incorrect (Figure 2C, grey histograms). We can demonstrate that the initial opinions are, on average, less biased when the correlated cue is incorrect, compared to when the correlated cue is correct: the left mode is closer than the right mode to 1/2 if: (*pr_I_* − 1/2)^2^ < ((*pr_I_* + (1 − *p*)) − 1/2)^2^ = ((*pr_I_* − 1/2) + (1 − *p*))^2^; this is easily shown to be true if *r_I_* > 1/2, which is what we assume in our model. Because of this, stalemates are more likely to occur when the correlated cue is incorrect, while consensus more likely occurs when the correlated cue is correct. Therefore, stalemates effectively reject the correlated cue when it is incorrect, which serves to break the dominance of the correlated cue and restore the wisdom of crowds in complex environments.

### Spatial clustering and sparseness improves the filtering of inaccurate decisions

In general, we find that the higher the probability of a stalemate, the more accurate the collective decision if consensus is reached, across all parameter space, in both simple and complex environments. Therefore, to understand how different properties of the network affect collective accuracy, we need only to examine how these properties affect the probability of stalemates.

We examine the role of four network properties on the probability of stalemates: the group size (*N*), the rewiring probability (*β*), the node degree *k* (here expressed as the normalized degree, *k/N*), and the randomness of the distribution of initial opinions in the network (where a value of 0 indicates that like opinions are maximally clustered in the network, and a value of 1 indicates that initial opinions are randomly distributed in the network). We find that the probability of stalemates tend to increase as group size increases, as the interaction network becomes more clustered, as the number of neighbors decreases, and as the initial opinions are increasingly clustered (Figure 3).

**Figure 3.**
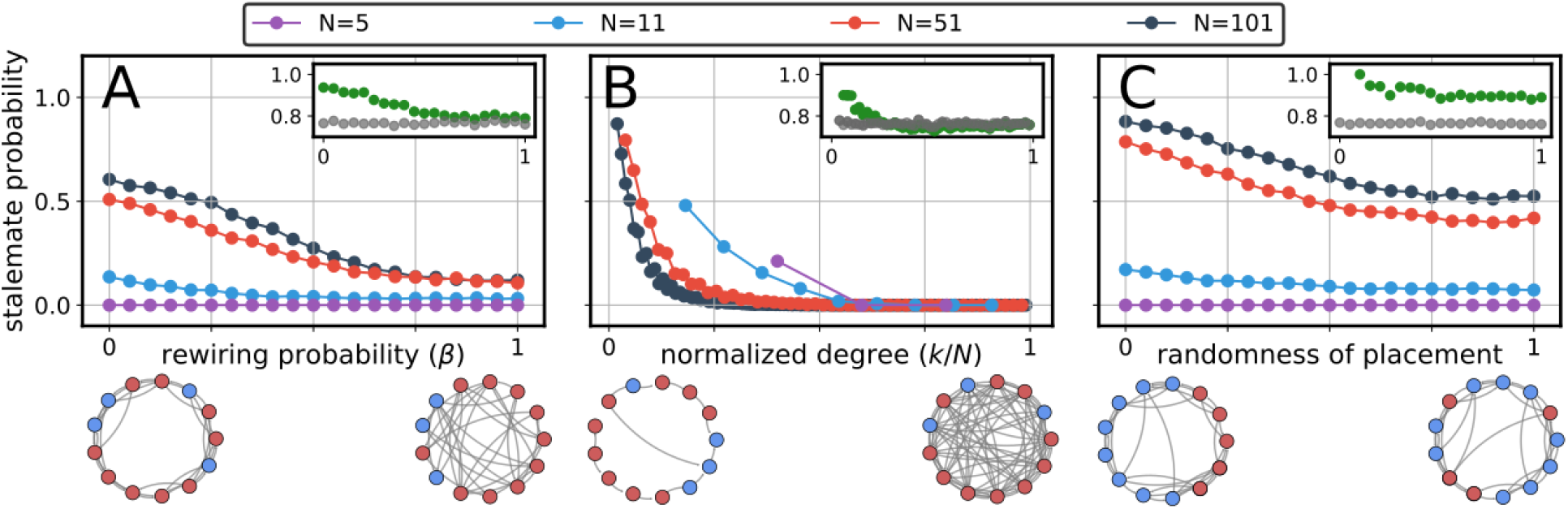
The effect of network structure on the probability of stalemates. These results are averages over all possible initial opinion configurations. We generally assume a default of *β* = 0.2, *k* = 5 and initial opinions that are randomly distributed within the network. Each panel shows the effect of varying a particular parameter while keeping the other two fixed. The insets show the effect of the respective structural parameters on collective accuracy for a group of *N* = 51 individuals in a simple environment with a cue of reliability *r* = 0.55. Green dots show collective accuracy at consensus and grey dots the result of a majority vote (unaffected by the structural parameters). In all three cases we find high values of collective accuracy to be linked to high probability of stalemate. **A.** The probability of stalemates increases as the rewiring probability shrinks (*i.e*., the network becomes more clustered structurally). This is particularly true for larger networks. **B.** The probability of stalemates increases as the average number of neighbors that an individual is connected to decreases (*i.e*., the network becomes sparser). **C.** The probability of stalemates increases as the distribution of initial opinions becomes more clustered. When the randomness parameter is 0, all nodes with the same opinion are placed next to each other in the network, and when the parameter is 1, initial opinions are randomly placed on the network.

Overall, then, the networks that lead to the highest probability of stalemate, and therefore the highest collective accuracy when consensus is reached, are those that are large, highly clustered (both in terms of their interaction structure and the distribution of initial opinions), and sparse (where individuals are each connected to few neighbors). Such social structures may in fact be common in nature, where visual occlusion [30, 31] or other mechanisms generate clustering in animal groups [22]. Furthermore, existing empirical evidence suggests that, for many social animal species, individuals pay attention to their closest 1-7 nearest neighbors [17, 18, 27, 32], resulting in relatively sparse social networks.

These findings have two main consequences, one descriptive and one prescriptive. The descriptive consequence is that, knowing the network structure and the specific decision context, one can predict the likelihood that a group reaches consensus through opinion dynamics. In general, there is a trade-off between the probability of consensus, and the collective accuracy when consensus is reached. The prescriptive consequence is that, since there is a trade-off between the probability of consensus and the collective accuracy when consensus is reached, individuals in the group may tune different structural parameters in order to titrate between consensus probability and decision accuracy.

### Faster decision bouts lead to increased collective accuracy

In addition to tuning the network’s structural parameters, we can also control parameters external to the network, such as the number of individual opinion updates (*s_max_*) permitted before a trial ends. Here again, we find that the harder consensus is to reach, the more likely it is to be accurate when reached (Figure 4). The reasoning is similar to that for clustered and sparse networks: in order for groups (particularly large ones) to reach consensus rapidly (*i.e*., when *s_max_* is small), the set of initial opinions must be highly biased, which in the case of positively informative cues tends to occur more often for the correct option.

**Figure 4.**
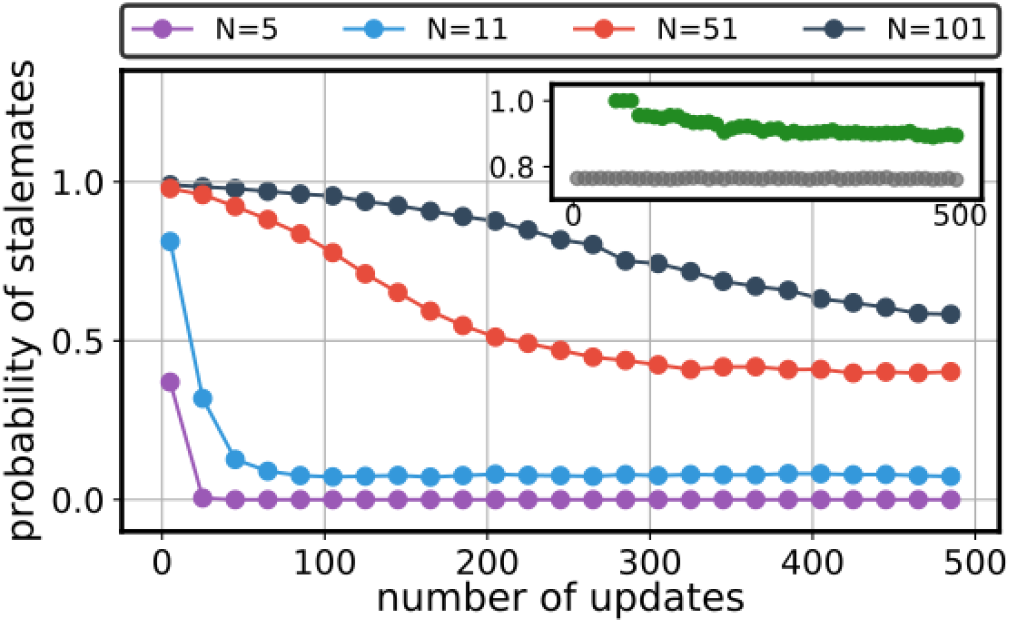
Probability of stalemates as a function of the maximum number of updating steps (*s_max_*) for groups of different sizes (*N* = 5 to 101). Small groups will almost always reach consensus after as few as 50 opinion changes. Larger collectives generally require more interactions to reach consensus and therefore only groups which are highly biased initially will manage to do so when interaction time is limited. Stalemate filtering leads to a decrease in collective accuracy as interaction time increases. As in Figure 3 insets show the effect of *s_max_* on collective accuracy for a group of *N* = 51 individuals in a simple environment with a cue of reliability *r* = 0.55. Green dots show collective accuracy at consensus and grey dots the result of a majority vote.

This result is seemingly in conflict with the well-known speed-accuracy trade-off, whereby higher decision accuracy typically comes with the cost of a greater amount of time needed to accumulate information and make a decision [33, 34]. In these cases, the trade-off is a direct consequence of the decision threshold, which determines the amount of evidence necessary for a decision to be made. If the threshold is high, decisions require large amounts of evidence, leading to longer integration time but also robustness against noise and therefore higher accuracy. By contrast, lower thresholds require less evidence and allow faster decisions. These fast decisions are more susceptible, however, to random fluctuations and are therefore less accurate.

In our model, the speed-accuracy trade-off appears to be violated *within* a given decision bout, since in general, faster decisions (smaller *s_max_*) lead to higher accuracy (Figure 4). However, this trade-off is restored when viewed *across* multiple decision bouts. When *s_max_* is small, each decision bout is short, but the number of bouts needed before an initial condition is found that is sufficiently biased to reach consensus increases. From this perspective, the total amount of time needed before a consensus decision is made increases as *s_max_* decreases, while the decision accuracy at consensus also increases – restoring the speed-accuracy trade-off.

### The reward structure modulates the optimal collective decision-making process

We have found that the probability of stalemates, and the collective accuracy when consensus is reached, can be adjusted by changing the size of the group, the number of neighbors that an individual is connected to, and how clustered the network and the distribution of information are. Many of these properties may plausibly be tuned either through evolution or learning. However, the optimal configuration of parameters will depend on the relative costs and benefits of stalemates, correct decisions, and incorrect decisions. So far, we have focused on decision accuracy when consensus is reached, implicitly assuming that stalemates are relatively cost-free, compared to incorrect decisions. Most models of collective decision making, by contrast, do not allow for stalemates, therefore assuming that stalemates are highly costly. The true cost of stalemates will be context dependent. For some contexts, such as a group of animals hiding in a shelter, the cost of stalemates will be low relative to the potential cost of leaving the shelter (and potentially being eaten by a predator). For other contexts, stalemates may be highly costly, such as when a predator is attacking a group and fleeing in almost any direction may be better than remaining in place due to indecision.

The interaction between the reward structure, network properties, and environmental properties is complex and beyond the scope of the current work, but here we simply illustrate how the reward structure can substantially alter the optimal structural parameters of the group. Figure 5 illustrates the effect of the three structural parameters (group size, rewiring probability, and network density) on a group’s relative reward, in both simple and complex environments. We examine three different reward structures (which are unitless). In the first scenario incorrect decisions are moderately more costly than stalemates. In this case we find that structural parameters associated with low stalemate probability are preferred in the simple environment while in the complex environment we find optimal intermediate values (Figure 5, second row). In the second scenario incorrect decisions are much more costly compared to stalemates. In simple environments this leads to optimal intermediate values of the structural parameters indicating that some nonzero probability of stalemates is preferable. In complex environments we find that structural parameters should be chosen such that the probability of a stalemate is maximal (Figure 5, third row). In the third scenario, incorrect decisions are again moderately more costly than stalemates, but both are more costly than correct decisions compared to the first scenario. Here in both environments the optimal choice is to choose structural parameters that avoid stalemates altogether (Figure 5, fourth row).

**Figure 5.**
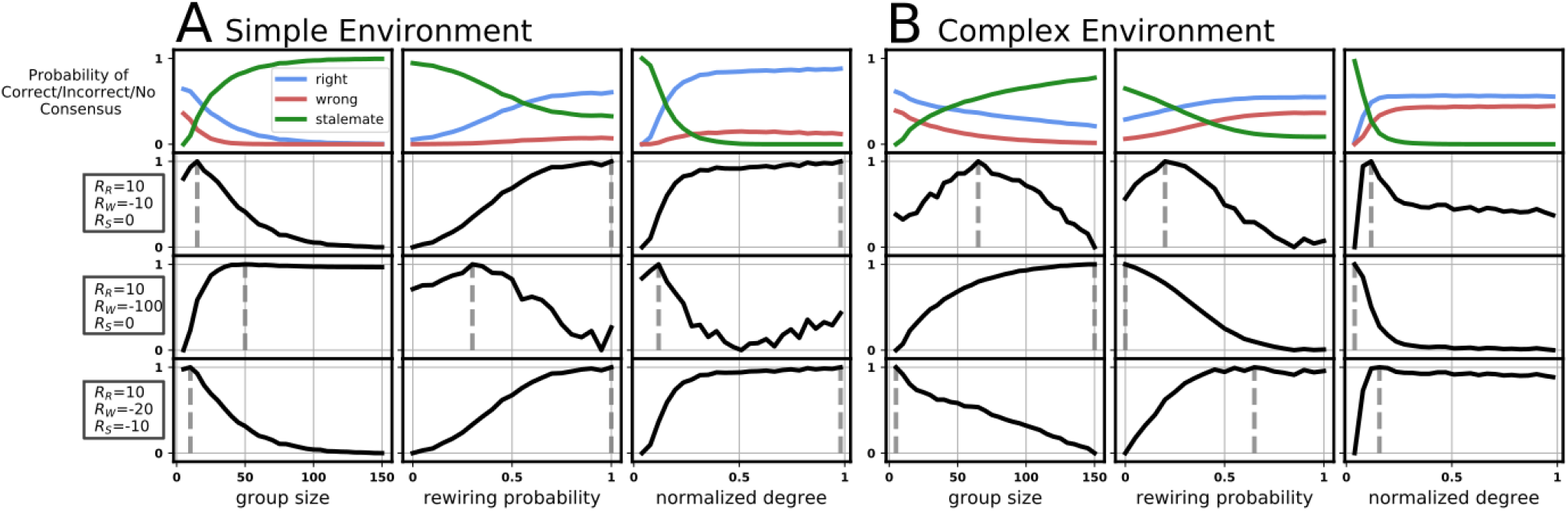
Relative reward as a function of different network structural parameters (group size *N*, rewiring probability *β*, and the normalized degree *k/N*) for simple (A) and complex (B) environments. As before, when varying one of the structural parameters, we keep the other two fixed at the default values (*N* = 51, *β* = 0.2 and *k* = 5 (*k/N* ≈ 0.098). For the simple environment, we set *r* = 0.58, and in the complex environment we set *r_I_* = 0.6, *r_C_* = 0.55, and *p* = 0.6, such that in both environments, a solitary individual makes a correct decision with probability 0.58. First row: the probability that a group makes a correct decision (blue), incorrect decision (red), or reaches a stalemate (green). Second row: a reward scenario where incorrect decisions have cost −10 (*R_W_*) and stalemates have cost 0 (*R_S_*). In all reward structures, correct decisions are associated with reward +10. In simple environments (A) this leads to a optimal parameters associated with low stalemate probability, while in complex environments (B) the optimal parameters are those associated with an intermediate probability of stalemates. Third row: incorrect decisions have cost −100, while stalemates have cost 0. In A this leads to optimal parameters associated with intermediate probability of stalemates, while in B the optimal parameters leads to a high rate of stalemates. Fourth row: incorrect decisions have cost −20, while stalemates have cost −10. In both A and B, the optimal parameters lead to a low rate of stalemates.

There is therefore a complex relationship between the relative rewards given by each potential outcome, and the optimal structural parameters that maximize expected reward. However, there may exist simple heuristics that may allow animals in groups to titrate their social network across contexts to improve, if not maximize, their expected rewards.

## Discussion

Our results highlight the importance of considering social decision-making dynamics that do not impose consensus and cannot be summarized simply as a form of ‘majority rule.’ In relaxing this assumption, we observe that such dynamics can often result in a stalemate, whereby the group is unable to reach a consensus. In our model, stalemates tend to filter outcomes where the majority decision would be incorrect, both in simple and complex environments, resulting in an improvement in collective accuracy.

For animal groups, using stalemates to filter out incorrect decisions would improve the fitness of the individuals in the group if the cost of indecision is low, relative to the cost of an incorrect decision. This is likely to be the case across a broad range of behaviorally and ecologically relevant contexts. Staying put may be preferable to misjudging the presence of a predator, getting lost, or moving to a lower-quality foraging patch. However, for other species or contexts, the cost of indecision is not low. For example, if a predator is attacking the group, when a food patch becomes completely depleted, or when a shelter becomes uninhabitable, indecision may be costly, and making even a wrong decision may be preferable to a stalemate. Therefore, the particular cost-reward structure of the options available to an animal group may either incentivize, or deincentivize, groups to employ stalemates as part of their decision-making process.

Individual animals in a group may, in principle, be able to titrate the frequency of stalemates by changing the proportion of long-distance connections, or the number of neighbors to which they’re attending. In principle, such changes to network structure can be achieved by even minor changes to individual behavioral rules. For example, fish schools and other social animal groups have long been known to move closer together in response to increased predation risk. A dominant early explanation for this tendency was that the close proximity resulted from competition for low-risk places within the group [2]. More recent work has demonstrated, however, that moving closer alters the network of social interactions in a way that increases collective responsiveness [35, 36, 37]. Highly responsive dense groups are characterized by social networks with fewer neighbors and higher clustering [30, 31]. In the context of our results, behavior such as moving closer would tend to increase the probability of stalemates, which may allow a group to stay in a shelter until they are certain that they are making a correct decision.

While many existing models, and experiments, focus on binary (two-choice) scenarios for simplicity, real world scenarios generally include at least one other possibility: indecision. Binary choice scenarios obscure the quantification of the costs and rewards underlying the two options, since the only relevant calculation is simply which option leads to a greater reward than the other. With three or more options (including indecision), the costs and rewards of each option must be made explicit to be able to order them. With the costs and rewards of the options made explicit, it may become clearer to what extent stalemates may be adaptive to a particular social species and context. For an animal group unable to choose a direction in which to flee, or a group stuck in a shelter, however safe, stalemates will be maladaptive. On the other hand, stalemates can cause a group to gather more information before making a decision. This can allow a group to avoid costly mistakes and boost collective accuracy, and occasional stalemates may be evolutionarily favored if the cost of indecision is relatively low to individuals in the group.

## Acknowledgements

CW, JBC and PR acknowledge the support of the *Cooperation and Collective Cognition Network (CoCCoN)* funded by the strategic partnership between Princeton University and Humboldt Universität zu Berlin. ABK acknowledges support from a Baird Scholarship and an Omidyar Fellowship from the Santa Fe Institute. PR acknowledges funding by the Deutsche Forschungsgemeinschaft (DFG, German Research Foundation) under Germany’s Excellence Strategy – EXC 2002/1 “Science of Intelligence” – project number 390523135, as well as through the Emmy Noether program, project number RO4766/2-1.

## Authors’ contributions

PR and JBC conceived the study. CW and PR designed the study with the help of ABK and JBC. CW implemented the model and performed numerical simulations and mathematical analyses. CW analyzed the simulation data with the help of PR, ABK and JBC. ABK wrote the paper with help from all authors. PR, CW, JBC critically revised the manuscript. All authors gave final approval for publication and agree to be held accountable for the work performed therein.

